# A Tail Fiber Protein and a Receptor-Binding Protein Mediate ICP2 Bacteriophage Interactions with *Vibrio cholerae* OmpU

**DOI:** 10.1101/2021.03.08.434494

**Authors:** Andrea N.W. Lim, Minmin Yen, Kimberley D. Seed, David W. Lazinski, Andrew Camilli

**Affiliations:** Department of Molecular Biology and Microbiology, Tufts University, School of Medicine, Boston, Massachusetts, USA; Department of Molecular Biology and Microbiology, Graduate School of Biomedical Sciences, Tufts University, School of Medicine, Boston, Massachusetts, USA

## Abstract

ICP2 is a virulent bacteriophage (phage) that preys on *Vibrio cholerae*. ICP2 was first isolated from cholera patient stool samples. Some of these stools also contained ICP2-resistant isogenic *V. cholerae* strains harboring missense mutations in the trimeric outer membrane porin protein OmpU, identifying it as the ICP2 receptor. In this study, we identify the ICP2 proteins that mediate interactions with OmpU by selecting for ICP2 host-range mutants within infant rabbits infected with a mixture of wild type and OmpU mutant strains. ICP2 host-range mutants had missense mutations in putative tail fiber gene *gp25* and putative adhesin *gp23*. Using site-specific mutagenesis we show that single or double mutations in *gp25* are sufficient to generate the host-range mutant phenotype. However, at least one additional mutation in *gp23* is required for robust plaque formation on specific OmpU mutants. Mutations in *gp23* alone were insufficient to give a host-range mutant phenotype. All ICP2 host-range mutants retained the ability to plaque on wild type *V. cholerae* cells. The strength of binding of host-range mutants to *V. cholerae* correlated with plaque morphology, indicating that the selected mutations in *gp25* and *gp23* restore molecular interactions with the receptor. We propose that ICP2 host-range mutants evolve by a two-step process where, first, *gp25* mutations are selected for their broad host-range, albeit accompanied by low level phage adsorption. Subsequent selection occurs for *gp23* mutations that further increase productive binding to specific OmpU alleles, allowing for near wild type efficiencies of adsorption and subsequent phage multiplication.

**Importance:** Concern over multidrug-resistant bacterial pathogens, including *Vibrio cholerae*, has led to a renewed interest in phage biology and their potential for phage therapy. ICP2 is a genetically unique virulent phage isolated from cholera patient stool samples. It is also one of three phages in a prophylactic cocktail shown to be effective in animal models of infection and the only one of the three that requires a protein receptor (OmpU). This study identifies a ICP2 tail fiber and a receptor binding protein and examines how ICP2 responds to the selective pressures of phage-resistant OmpU mutants. We found that this particular co-evolutionary arms race presents fitness costs to both ICP2 and *V. cholerae*.

## Introduction

Vibriophages, phages that prey on bacteria from the gram-negative *Vibrio* genus, were first described by E.H. Hankin in 1896 as antimicrobial agents from the Ganges River, two decades before phages were formally identified by Twort and d’Herelle (1, 2). Growing concern over the emergence of multidrug-resistant strains of *Vibrio cholerae* (3), the causative agent of cholera, has renewed scientific interest in vibriophages as therapeutics (2, 4) and environmental markers of cholera outbreaks (5-7). In 2011, our lab isolated three unique virulent vibriophages from the rice-water stools of Bangladeshi cholera patients, designated phages ICP1, ICP2, and ICP3 (7). ICP2 is morphologically categorized as a short-tailed podovirus but bears little genetic homology to the canonical podovirus T7 or other members of the family. Its 50 kb genome encodes 73 putative protein coding sequences divided into two transcriptional units in opposite orientations.

An isolate of ICP2 was also recovered in 2014 from the rice-water stool of a Haitian cholera patient (8). This sample also contained several ICP2-resistant isogenic *V. cholerae* strains that had missense mutations in *ompU*. OmpU is an outer membrane general porin that is associated with virulence (9-11), adherence (12), and osmoregulation (13-15). ICP2-resistant strains with null mutations in *toxR* were also found among several stool samples containing ICP2. ToxR is a transcriptional activator of virulence genes including *ompU*. Accordingly, the *toxR* null mutations resulted in attenuated *V. cholerae* colonization in an infant mouse model as well as the complete loss of *ompU* expression (9). Further experiments showed that two OmpU missense mutations, V324F and G325D, only minimally affect *V. cholerae* fitness. These mutations identified OmpU as the ICP2 receptor and demonstrated how vibriophages impose selective pressures during cholera infections.

In 2018, two groups solved the crystal structure of *V. cholerae* OmpU (16, 17). OmpU is an outer membrane localized homotrimer, with each monomer comprised of a 16-β-stranded barrel. Each monomer has a central pore of ∼0.55-0.6 nm wide (17) and has 8 protruding extracellular loops (Figure 1). L2 projects into the neighboring monomer, while L3 is a constriction loop that bends into the center of each barrel. The OmpU residues that mutated to confer ICP2 resistance in the stool samples (8) are highlighted on one monomer-monomer interface in Figure 1 (8, 16). All mutations, except N158Y, are in L4 and L8. These extracellular loops are adjacent between neighboring monomers. N158Y is found near the transmembrane region of L3, and its affect on OmpU localization or structure are unknown. These observations suggest that ICP2 binds *V. cholerae* at the L4 and L8 interfaces of OmpU trimers. We sought to identify the ICP2 proteins that interact with OmpU and examine how ICP2 may counter-evolve in response to phage-resistant *V. cholerae* OmpU mutants.

**Figure 1.**
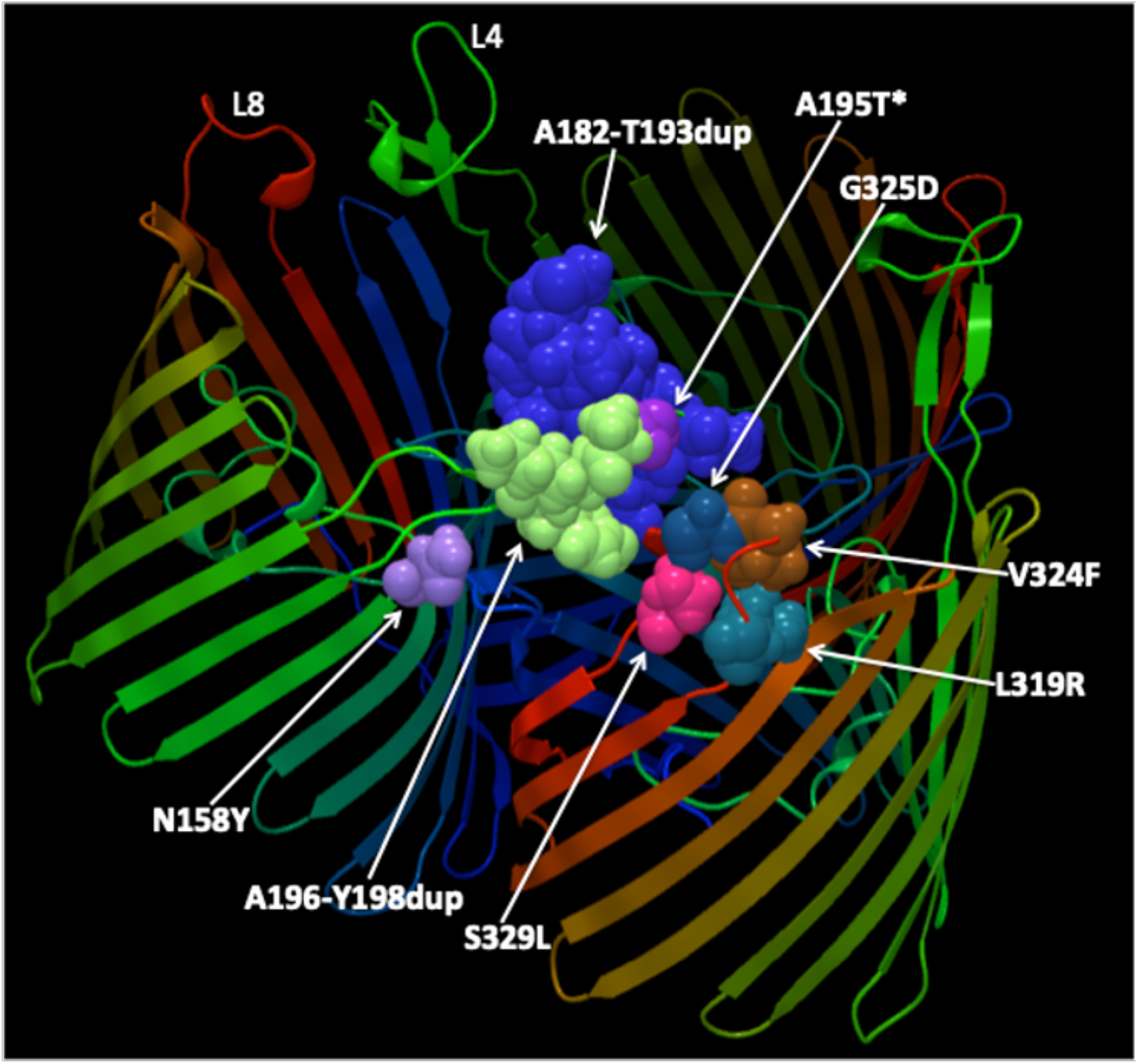
Location of mutated amino acid residues in the OmpU structure that give rise to ICP2-resistance. OmpU mutations previously shown to confer ICP2 resistance (8) map to extracellular loops L4 and L8 on the structure solved by Li et al. (16). The OmpU porin is a homotrimer of three β-barrels. Mutations are highlighted on two adjacent monomers to emphasize their proximity. *Mutation A195T only confers partial resistance to ICP2 (8). This image does not depict how these mutations may affect OmpU structure.

## Results

### ICP2 host-range mutants have missense mutations in *gp23* and *gp25*

ICP2 host-range mutants were selected during intestinal infection of infant rabbits co-inoculated with ICP2 and a mixture of *V. cholerae* wild type (WT) and OmpU point mutant strains at a 10:1 ratio. The addition of WT *V. cholerae* was designed to allow for enough replication of ICP2 to yield spontaneous host-range mutants that could then be enriched by replicating on the OmpU mutants. Until the appearance of such host-range mutants, the OmpU mutants have a competitive advantage over WT *V. cholerae* due to lack of predation by ICP2. Indeed, this was shown previously when ICP2 predation resulted in a 10,000-fold competitive advantage for OmpU G325D over the WT after 12 hours of infection of infant rabbits (8). One group of four animals was inoculated with WT *V. cholerae* and a mixture of OmpU A196_Y198dup, OmpU A195T, OmpU L319R, and OmpU S329L. A second group of four animals was inoculated with WT and a mixture of OmpU A182_T193dup, OmpU V324F, and OmpU N158Y. A third group of four animals was infected with WT and OmpU G325D. ICP2 in cecal fluid obtained from euthanized symptomatic animals was plaqued on soft agarose overlays containing specific OmpU mutants to isolate host-range mutants. These included OmpU mutants A195T, V324F, and G325D. Up to seven plaques per host strain per animal were chosen for plaque purification on each respective OmpU mutant. All plaques were clear and of equivalent size as those formed by WT ICP2 on WT *V. cholerae*.

The ICP2 host-range mutants were whole-genome sequenced to identify possible mutations. Of the 28 isolates sequenced, thirteen were unique (Figure 2A). Those isolated on OmpU V324F or OmpU G325D contained multiple single nucleotide variants (SNVs) in *gp23* and *gp25* (Figure 2), each a missense mutation. This strongly implicates Gp23 and/or Gp25 as ICP2 proteins that interact with the OmpU receptor. According to homology analysis via Interative Threading Assembly Refinement (I-TASSER) (18-20) and the Rapid Annotation using Subsystem Technology version 2 (RAST) (21, 22), *gp23* encodes a putative receptor binding adhesin protein and *gp25* encodes a putative phage tail fiber. Although *gp24* is also annotated as a putative tail fiber, no mutations were found in this gene. It is unlikely that Gp23 and Gp25 are tail tube proteins because those have previously been identified bioinformatically as Gp9, Gp10, and Gp11 (23).

**Figure 2.**
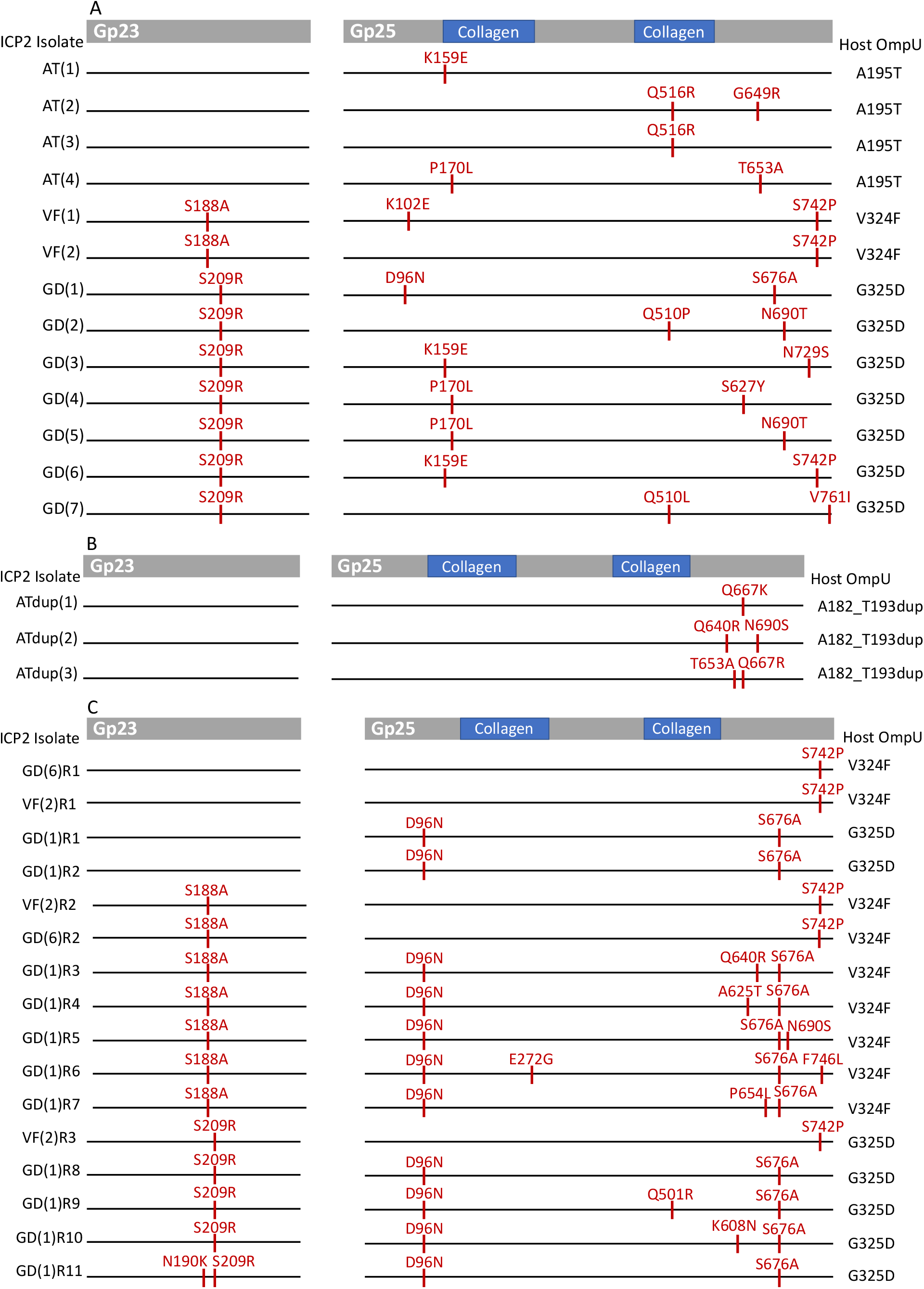
ICP2 host-range mutants have missense mutations in *gp23* and *gp25*. Thick grey bars represent the WT amino acid sequence of Gp23 and Gp25, with putative domains indicated where possible. Each pair of thin horizontal lines represents the Gp23 and Gp25 of a single ICP2 mutant (denoted on the left). The mutations present in each isolate are above their corresponding red tick marks. The OmpU allele that each phage mutant was isolated on is indicated on the right. **(A)** ICP2 host-range mutants isolated from the cecal fluid of infant rabbit co-infections had mutations in *gp23* and *gp25*. **(B)** ICP2 host-range mutants selected *in vitro* on OmpU A182_T193dup had mutations in *gp25*. **(C)** Recombinant ICP2 host-range mutants were generated *in vitro* through sequential infections of WT *V. cholerae* strains carrying plasmids with specific *gp23* or *gp25* mutations. Mutants were selected based on their ability to form plaques on either OmpU V324F or OmpU G325D. Gp25 mutations were derived from VF(2), GD(1), and GD(6). Either Gp23 S188A or S209R was then added to these mutants. Several isolates derived from GD(1) acquired one or two novel mutations during the second recombination infection.

ICP2 mutants isolated on OmpU V324F have a Gp23 S188A mutation, while those isolated on OmpU G325D have a Gp23 S209R mutation (Figure 2). Gp25 mutations were more prevalent and varied among the ICP2 host-range mutants. Gp25 mutations clustered near the N- and C-termini, which are outside or near the ends of two predicted collagen fiber-like domains (Figure 2). Gp25 mutations exhibit loose specificity for individual OmpU alleles. Residues Q510 and N690 both mutated to different amino acids in different ICP2 phages isolated on the OmpU G325D mutant. In contrast, a single S742P mutation was shared among phages isolated on both OmpU V324F and G325D mutants. Because these ICP2 host-range mutants were selected during intestinal infection, we could not determine how many rounds of phage replication occured before developing the ability to infect an OmpU mutant nor the order in which Gp23 and Gp25 mutations were selected. Therefore, we next sought to parse the relationship between the ICP2 Gp23 and Gp25 mutations and *V. cholerae* OmpU alleles.

### ICP2 host-range mutants require at least one Gp25 mutation

Plasmid-based recombination in *V. cholerae* was used to generate ICP2 mutants with the *gp25* mutations identified in VF(2), GD(6), and GD(1). Host-range mutants are herein named for the OmpU mutant host on which they were isolated followed by a unique identifying number. VF(2) and GD(6) both contain a Gp25 S742P mutation despite being isolated on different OmpU mutants. Gp25 in GD(6) also contains a unique second K159E mutation. GD(1) contains Gp25 D96N and S676A mutations. Recombinant plaques with single Gp25 S742P mutations, derived from VF(2) or GD(6), were isolated and plaque-purified once on OmpU V324F and then amplified on WT. GD(1) recombinants were isolated and plaque-purified once on OmpU G325D and then amplified on WT. We chose to amplify on WT instead of an OmpU mutant in order to limit further selection for spontaneous mutations that might increase infectivity on the latter host. All host-range mutants isolated in this study formed clear, normal sized plaques on WT, suggesting there would not be selective pressure for additional mutations on this host. Control recombination infections using a plasmid containing WT *gp23* and *gp25* did not result in any host-range mutants. The recombinant mutants were whole-genome sequenced to verify mutations. Recombinant mutants are designated with an “R” and a unique identifying number following their parent name (Figure 2C).

Efficiency-of-plaquing (EOP) assays were used to quantify how efficiently these mutants can infect OmpU V324F and G325D relative to WT cells. Low EOPs often correlated with a turbid plaque morphology (Figure 3A). Recombinant ICP2 mutants with only one or two Gp25 mutations maintain the ability to form clear plaques on WT host cells but can only weakly form plaques, i.e., low EOPs and turbid plaques, on either OmpU V324F or OmpU G325D. No host-range mutants formed plaques on a Δ*ompU* control, indicating that they are interacting with either OmpU V324F or G325D and not another receptor. Isolates VF(2)R1 and GD(6)R1 are biological duplicates with a Gp25 S742P mutation. While they form turbid plaques on both OmpU V324F and G325D, their EOP on OmpU V324F is only 10-fold lower than on WT (Figure 3A). In contrast, their EOPs on OmpU G325D are 4- to 5-logs lower than on WT. GD(1)R1 and GD(1)R2, with Gp25 mutations D96N and S676A, also form turbid plaques on both OmpU V324F and G325D. These phage mutants have a clear preference for OmpU G325D, with GD(1)R2 barely reaching a statistically significant EOP on OmpU V324F. While Gp25 mutations alone do not provide robust phenotypes, at least one is required to weakly infect an OmpU mutant.

**Figure 3.**
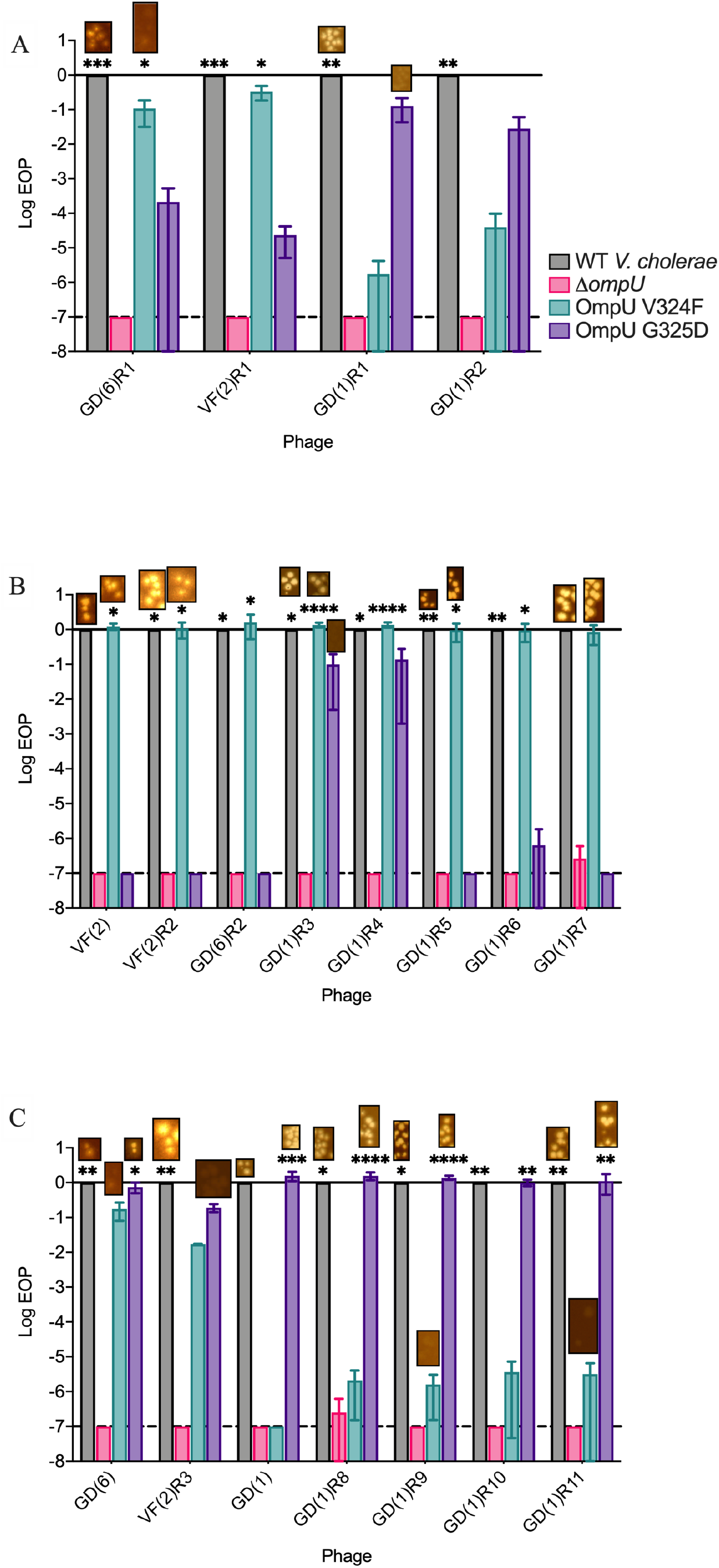
Efficiency of Plaquing of ICP2 host-range mutants on OmpU V324F and G325D. EOPs were determined relative to that on WT *V. cholerae*. EOP assays were done in at least quadruplicate with a starting titer of ∼10^8^ PFU/ml. The dotted line represents the limit of detection and the bars and vertical lines show mean and standard deviation. No data points were included for plaques that were too turbid to count. Examples of plaque morphology are shown above their associated bars. Plaque assays were scanned using Epson scan (V. 3.25A) with adjustments made to contrast and histogram input and output to best visualize turbid plaques. Scanner settings were kept the same for all plates in a single replicate. Statistical significance was determined relative to no plaque formation on Δ*ompU* (Ordinary one-way ANOVA and *post hoc* Dunnett’s multiple comparisons test on log-transformed data points). (* P ≤ 0.05, ** P ≤ 0.01, *** P ≤ 0.001, ***** P ≤ 0.0001) **(A)** Recombinant ICP2 mutants with only one or two Gp25 mutations form turbid plaques on OmpU V324F or OmpU G325D. They also retain the ability to form clear plaques on WT cells. Recombinant isolates GD(6)R1 and VF(2)R1 each have a Gp25 S742P mutation. Recombinant isolates GD(1)R1 and GD(1)R2 contain the two Gp25 mutations found in GD(1). **(B)** ICP2 host-range mutants with secondary Gp23 S188A mutations form clear plaques on OmpU V324F and have EOPs near 1, regardless of their Gp25 mutations. Recombinant isolates GD(1)R3 and GD(1)R4 retain the ability to weakly infect OmpU G325D. **(C)** ICP2 host-range mutants with secondary Gp23 S209R mutations have increased EOPs on OmpU G325D, regardless of their Gp25 mutations.

Further support for the important role of Gp25 in conferring a host-range phenotype came from host-range mutants evolved in animals and then isolated on OmpU A195T, a host strain only partially resistant to WT ICP2 (8). These phage mutants have one or two mutations in Gp25 and no mutations in Gp23 (Figure 2A). This provides further evidence that Gp25 mutations are sufficient to expand host-range.

In host-range assays that included five additional ICP2-resistant OmpU mutants, ICP2 mutants with only one or two Gp25 mutations and no Gp23 mutations exhibit a similar host-range pattern despite the diversity of their Gp25 mutations (Figure 4, top eight ICP2 mutants). In addition to forming clear plaques on WT *V. cholerae*, most of these host-range mutants have higher EOPs on OmpU L319R, S329L, and A182_T193dup. This suggests that initial Gp25 mutations are necessary and sufficient to expand host-range.

**Figure 4.**
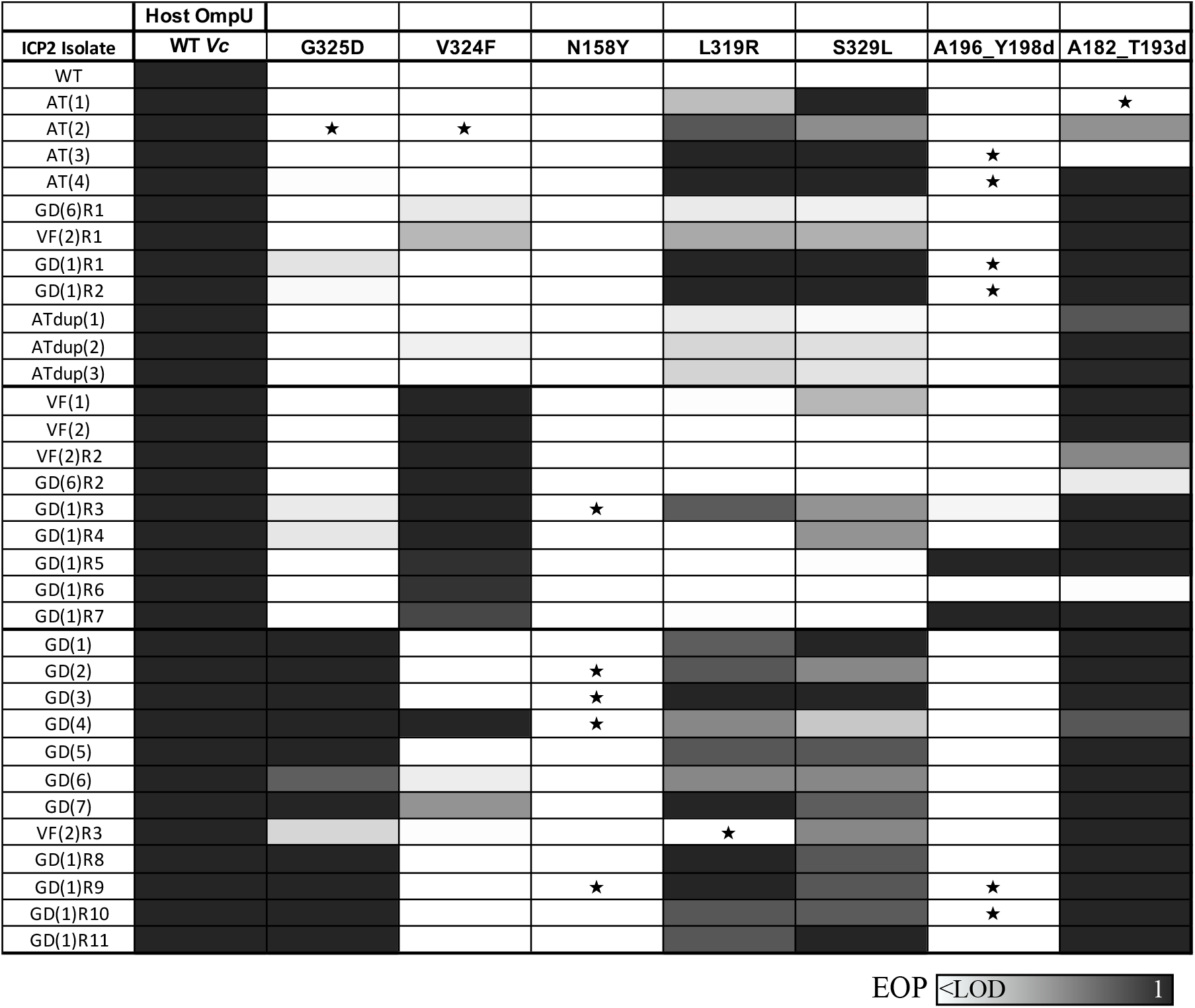
Host-range Efficiency of Plaquing assays on ICP2-resistent OmpU mutants. Host-range mutants are organized top to bottom rows according to their Gp23 mutations: the top 11, AT(1)-GD(1)R2, have no Gp23 mutation; the middle nine, VF(1)-GD(1)R7, have Gp23 S188A; the lower 11, GD(1)-GD(1)R10, have Gp23 S209R; and the bottom mutant, GD(1)R11, has Gp23 N190K and S209R. Approximate EOPs from below the limit if detection (<LOD) to 1, indicated by the scale-bar, are based on the mean of two to five replicates; averages >1 were set to 1. ICP2 mutants shown in Figure 3 were not retested on OmpU V324F and OmpU G325D; these boxes use the mean EOP values from Figure 3. ★ indicates that lysis at lower dilutions was observed in at least two replicates, without single plaques at higher dilutions.

### Selection of ICP2 host-range mutants *in vitro*

The observation that all ICP2 host-range mutants isolated in animals retain the ability to plaque on WT *V. cholerae* could be due to fact that both WT and OmpU mutants were present during the selection. To investigate this hypothesis, we sought to isolate host-range mutants *in vitro* on pure cultures of the OmpU A182_T193dup strain and then test plaque formation on WT *V. cholerae*. Initial attempts to select ICP2 host-range mutants *in vitro* failed, likely due to lack of a sufficiently large enough pool of input phage to harbor preexisting mutants. To overcome this limitation, we generated two independent high-titer stocks of ICP2 on a mutator strain of *V. cholerae* in which the mismatch repair gene *mutS* was deleted. Deletion of *mutS* was previously shown to increase the mutation rate of *V. cholerae* by 100-fold (24). Although it is not known if the host methyl-directed mismatch repair system functions with ICP1 DNA replication, it is notable that ICP1 encodes its own DNA adenine methyltransferase (Dam) (8). After two rounds of selection on the OmpU A182_T193dup strain, host-range mutants that form clear plaques were obtained. Whole-genome sequencing of four ICP2 host-range mutants from each independent selection yielded a total of four lineages; one from the first and three from the second selection. Previously observed and novel mutations in *gp25* were found, but no mutations in *gp23* were present (Figure 2B). Two mutants contained the previously observed single Gp25 S742P mutation. Only the three host-range mutants with novel Gp25 mutations are shown in Figure 2B and were further analyzed. These results are consistent with those obtained from the *in vivo* selected host-range mutants, in that mutations in Gp25 are sufficient to give a host-range phenotype on an OmpU A182_T193dup strain.

Next, we compared EOP and plaque phenotype on OmpU A182_T193dup and WT *V. cholerae*. The three host-range mutants formed clear plaques on OmpU A182_T193dup and, as observed for the *in vivo* selected host-range mutants, all retained clear plaque formation on WT *V. cholerae* (Figure 4, rows 9-11). Consistent with this, their EOPs on OmpU A182_T193dup and WT were at or near 1. These mutants formed turbid plaques on OmpU L319R and S329L, similar to the other Gp25-only mutants. Therefore, retention of infection of WT *V. cholerae* in ICP2 host-range mutants appears to be a general phenomenon, and not the result of selection in the presence of WT and an OmpU mutant.

### Secondary Gp23 mutations increase EOP on specific OmpU mutants

To attempt to isolate ICP2 host-range mutants with only Gp23 mutations, WT *V. cholerae* with plasmids containing *gp23* S188A or S209R alleles were infected with WT ICP2. However, no recombinant plaques could be isolated on OmpU V324F or G325D. When the population of progeny phages from these infections were sequenced ∼1% of the phage were found to have the intended mutations. Therefore, it appears that Gp23 mutations alone are insufficient to give a host-range phenotype.

A secondary Gp23 S188A in Gp25-only mutants imparts clear plaque formation on OmpU V324F. Host-range mutants with secondary Gp23 mutations were generated by using the ICP2 recombinants with one or two Gp25 mutations to infect WT cells containing plasmids containing either *gp23* S188A or S209R alleles. The addition of a Gp23 S188A leads to more efficient plaquing on OmpU V324F for all seven recombinants regardless of initial Gp25 mutations (Figure 3B). VF(2), VF(2)R2, and GD(6)R2 are biological triplicates containing Gp25 S742P and S188A, but were generated in an animal and *in vitro*, respectively. GD(1)R3, GD(1)R4, GD(1)R5, GD(1)R6, and GD(1)R7 contain Gp25 D96N and S676A from GD(1), as well as one to two novel Gp25 mutations (Figure 2C). Gp23 S188A was not seen with these combinations of Gp25 mutations among the host-range mutants evolved *in vivo*.

Similarly, a secondary Gp23 S209R mutation results in increased EOPs and/or clear plaque formation for all five recombinants on OmpU G325D regardless of initial Gp25 mutations (Figure 3C). VF(2)R3, containing Gp25 S742P and Gp23 S209R, was the only recombinant that still forms turbid plaques on OmpU G325D, but its EOP of 0.19 is markedly higher than that of Gp25 S742P mutant VF(2)R1, which has a mean EOP of 2.3 × 10^−5^. The additional Gp25 K159E mutation in GD(6) increased the EOP further and gave clear plaques on OmpU G325D. The presence of Gp25 S742P also likely allows both GD(6) and VF(2)R3 to form turbid plaques on OmpU V324F. GD(1)R9 and GD(1)R10 both have novel Gp25 mutations in addition to S676A and D96N mutations present in GD(1), while GD(1)R11 has a second Gp23 N190K mutation. These new mutations, however, do not result in EOPs on OmpU V324F and G325D that greatly differ from GD(1) or GD(1)R8 (Figure 3C). These results suggest that Gp23 mutations increase infection in a mostly allele-specific manner with respect to OmpU mutants, while retaining wild-type infection of WT OmpU.

In EOP assays that included five additional OmpU mutants, ICP2 mutants with secondary Gp23 mutations have modest alterations in host range (Figure 4). The middle 9 rows in Figure 4 show the host range of mutants with secondary Gp23 S188A mutations and the mutants in the final 12 rows all have a secondary Gp23 S209R mutation. The addition of a secondary Gp23 S188A mutation in VF(2), VF(2)R2, and GD(6)R2 corresponds with an inability to form plaques on OmpU L319R and S329L. VF(2)R2 and GD(6)R2 were derived from VF(2)R1 and GD(6)R1, which contain a single Gp25 S742P mutation and could previously infect these strains. VF(2)R3 contains the same single Gp25 S742P mutation and a secondary Gp23 S209R mutation but retains the ability to infect OmpU S329L and very weak infection of OmpU V324F. In addition to secondary Gp23 mutations, the presence of novel Gp25 missense mutations may also be playing a role in the host ranges of the remaining ICP2 mutants.

### ICP2 host-range mutants can bind to OmpU mutants

If plaque formation on OmpU V324F and G325D requires direct binding by Gp23 and/or Gp25, binding to these OmpU alleles should correlate with EOP. We assayed binding over a period of 24 hours (hrs) to heat-killed host cells and quantified binding as the ratio of remaining free phage titer to the initial titer at t = 0 hrs. Cells were heat-killed by incubating at 51°C for 12 minutes (min) in a PCR thermocycler. The structure of the cells was unaffected by this treatment as determined by phase-contract microscopy. WT ICP2 bound to WT *V. cholerae* between 10- and 100-fold, depending on the experiment, but did not exhibit any OmpU-independent binding to the *ΔompU* strain. ICP2 host-range mutants all retain the ability to significantly bind WT *V. cholerae* (Figure 5).

**Figure 5.**
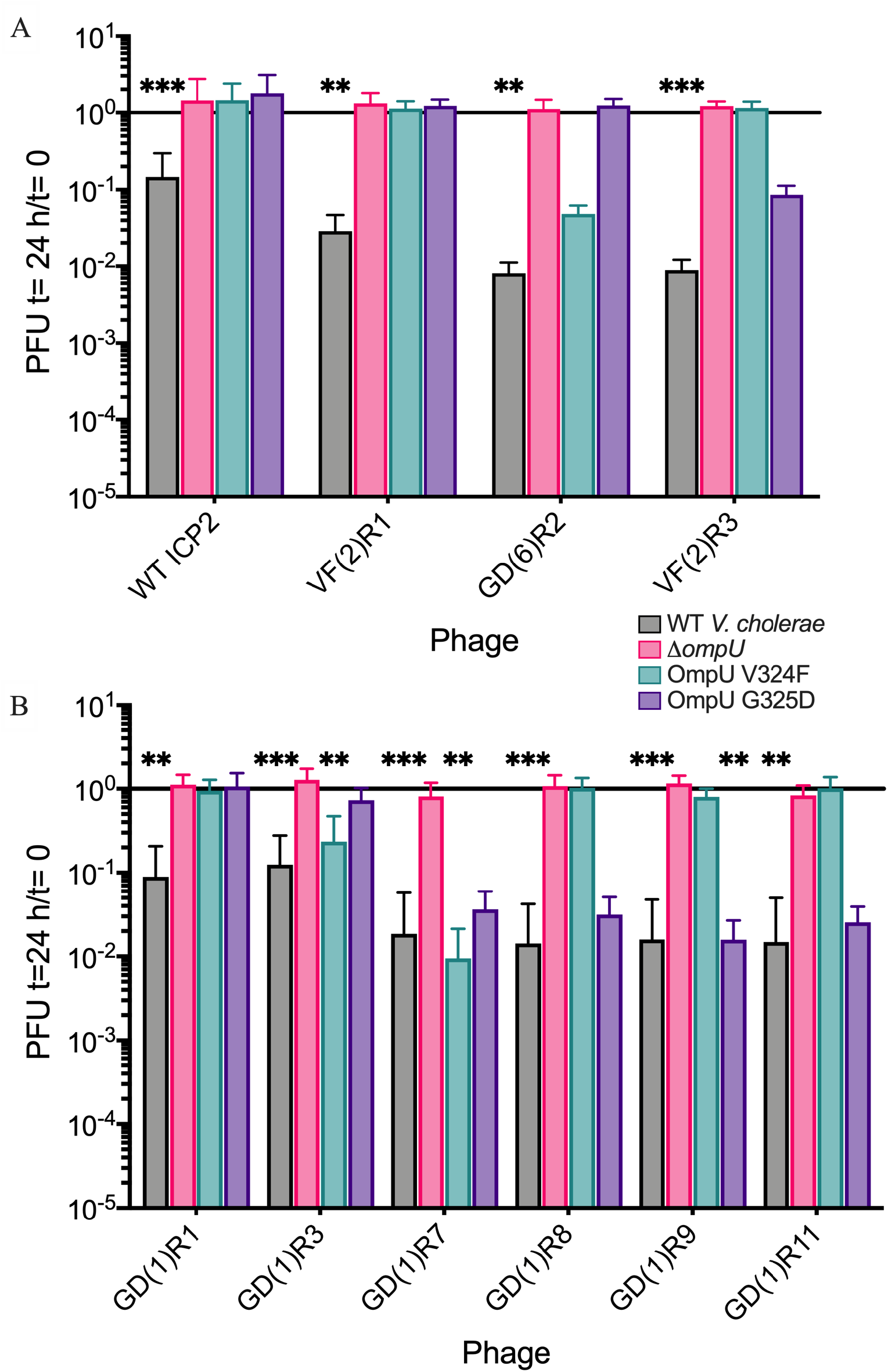
Phage binding to heat-killed OmpU mutant cells. Phage were added to heat-killed cells at an MOI ∼ 0.1 and incubated for 24 hrs at room-temperature. Binding was determined as the ratio between PFU added (t=0) to PFU remaining at t=24 h; a ratio near 1 indicates no detectable binding. Each bar represents the mean and standard deviation of four to 12 biological replicates (WT ICP2 n=12, GD(1)R11 on OmpU G325D n=4, most samples have six to nine biological replicates). Statistical significance was determined relative to no binding to Δ*ompU* (Kruskal-Wallis and *post hoc* Dunn’s multiple comparison tests). (* P ≤ 0.05, ** P ≤ 0.01, *** P ≤ 0.001, ***** P ≤ 0.0001). **(A)** WT ICP2 only binds to heat-killed WT *V. cholerae* cells. VF(2)R1 also only binds WT *V. cholerae* despite having a single Gp25 S742P mutation. The addition of Gp23 S188A or S209R is associated with binding to OmpU V324F or G325D, respectively. GD(6)R2 and VF(2)R3 bind OmpU V324F and G325D as expected, but not at statistically significant amounts. **(B)** Similarly, GD(1)R1 has two Gp25 mutations but only binds to WT *V. cholerae*. The addition of Gp23 S188A or S209R results in the ability to bind specific OmpU alleles. Binding to OmpU G325D does not reach statistical significance. GD(1)R7 binds both OmpU alleles despite only forming plaques on OmpU V324F.

VF(2)R1 and GD(1)R1 show no binding to OmpU V324F and G325D (Figure 5), respectively. Because these phage mutants could form turbid plaques and had low to intermediate EOPs on these hosts, this suggests that the binding assay is not as sensitive. ICP2 mutants with a Gp23 S188A mutation, GD(6)R2 (Figure 5A), GD(1)R3, and GD(1)R7 (Figure 5B), all bind to OmpU V324F, as expected considering these phage mutants have high EOPs and form clear plaques. Similarly, VF(2)R3 (Figure 5A), GD(1)R8, GD(1)R9, and GD(1)R11 (Figure 5B) contain Gp23 S209R and bind to OmpU G325D. The one exception to this trend is that GD(1)R7, despite not forming plaques or having a detectable EOP on OmpU G325D, binds to both OmpU alleles with a preference for OmpU V324F.

In general the host-range mutants bind better to WT cells than to OmpU V324F and G325D mutants. VF(2)R1 has no detectable binding to either OmpU V324F or G325D. GD(6)R2 and VF(2)R3 show 99% binding to WT *V. cholerae*, but 95% and 91% binding to OmpU V324F and G325D, respectively (Figure 5A). Host-range mutants with significant binding to OmpU V324F or G325D are derived from GD(1)R1 or GD(1)R2 and have an additional Gp25 mutation (Figures 2B, 5B). The additional Gp23 mutation in GD(1)R11 does not significantly aide binding to OmpU G325D.

### ICP2 host-range mutants prey on OmpU mutants in broth culture with varying efficiencies

The binding assays demonstrated that phage predation in broth culture differs from the slower diffusion of soft agar overlays in that the on-off rate of receptor binding will have a larger impact on the rate of adsorption. Therefore, we conducted predation assays in shaking broth cultures to examine how the ICP2 phenotypes seen in EOP and binding assays are reflected on host cell killing. The cell density of early exponential growth phase *V. cholerae* cultures infected with ICP2 host-range mutants (MOI ∼1) was measured as the optical density at 600 nm over 16 hrs at 37°C. As previously reported (8), the negative control *ΔompU* strain has a slight growth defect in LB (Figure 6A). All ICP2 host-range mutants prey on WT cells, suppressing exponential growth for ∼6 hrs, followed by a return to exponential growth (Figure 6). Most *V. cholerae* cells replicating after 6 hrs have gained phage resistance (Table S1).

**Figure 6.**
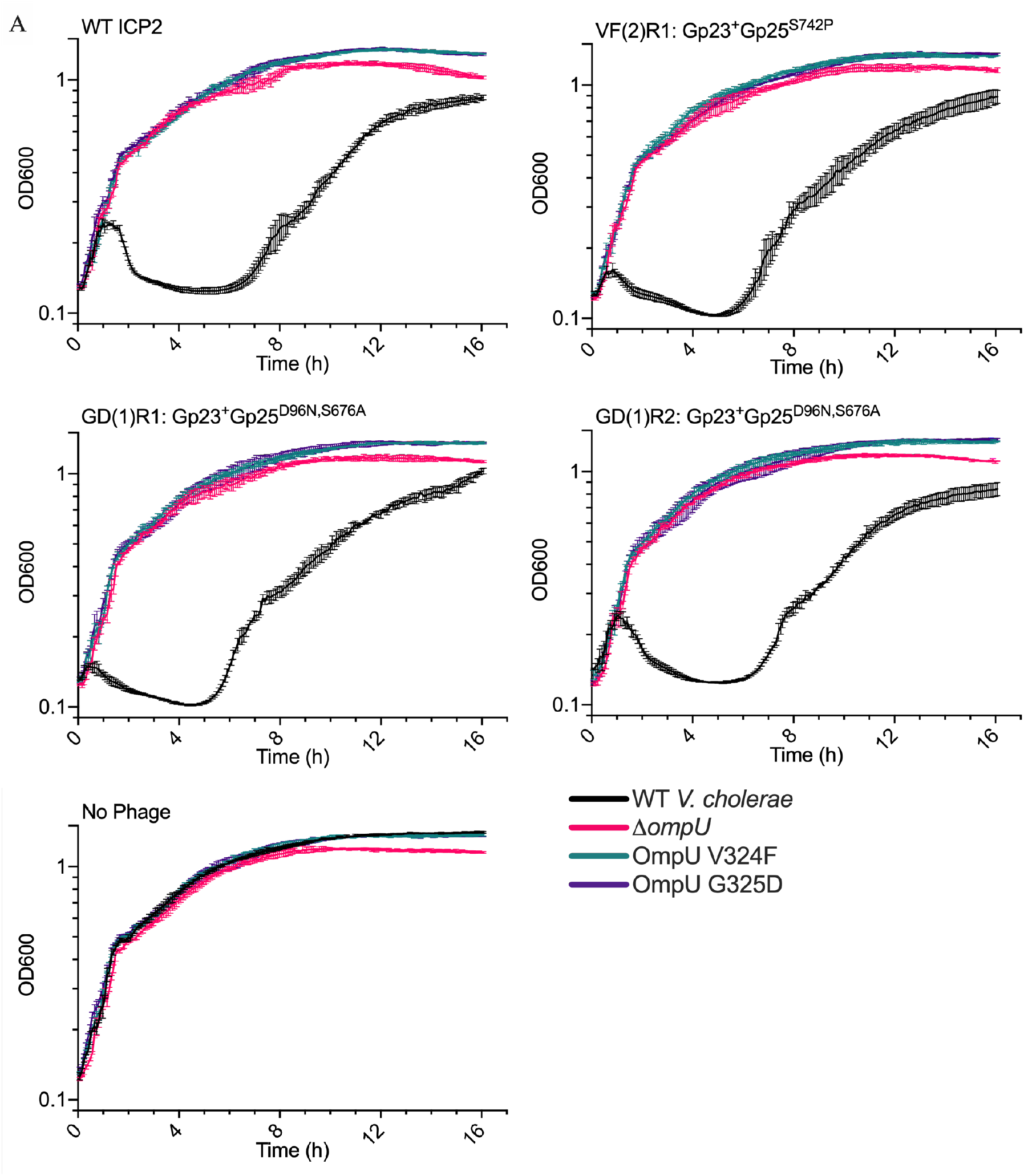
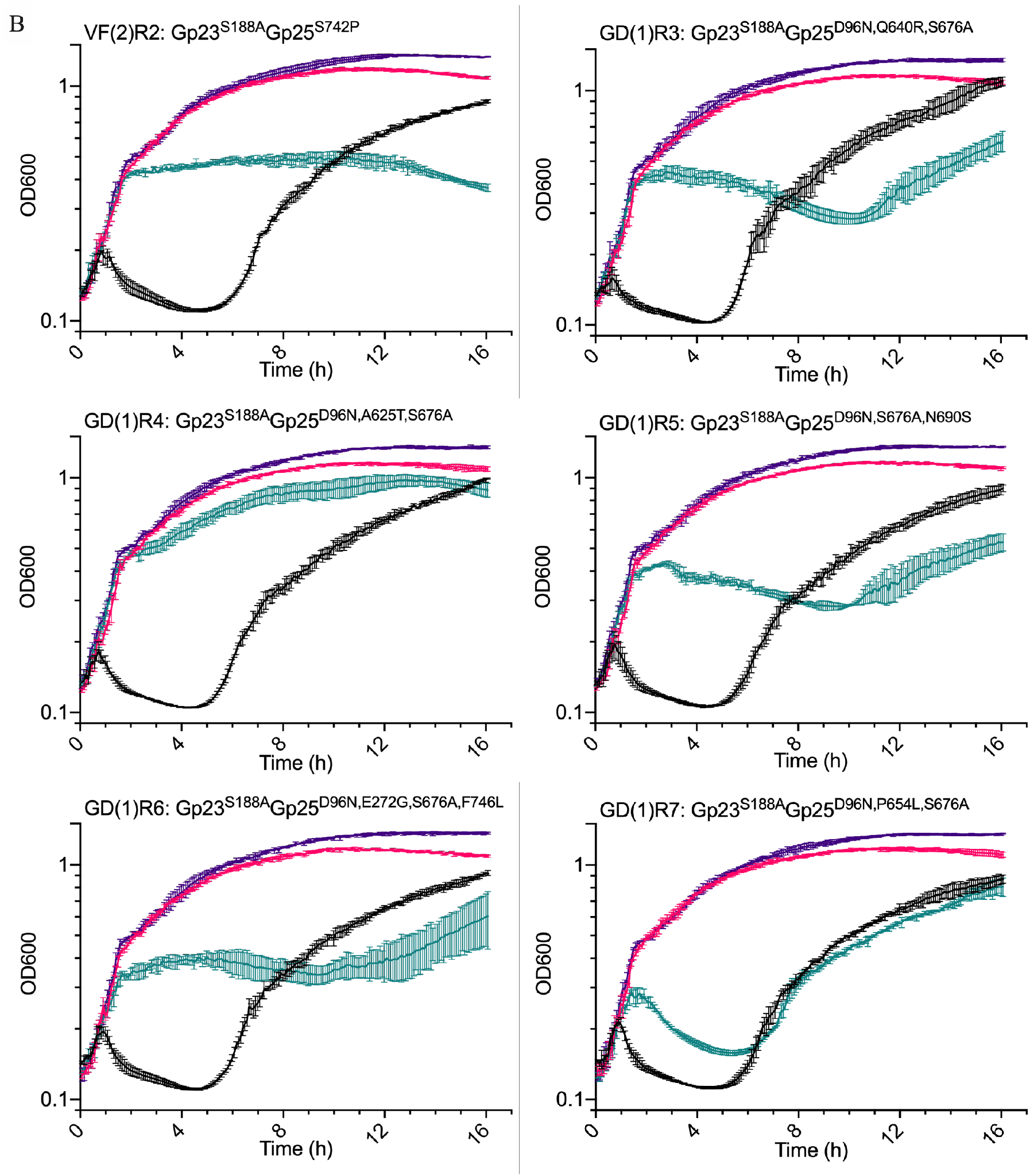
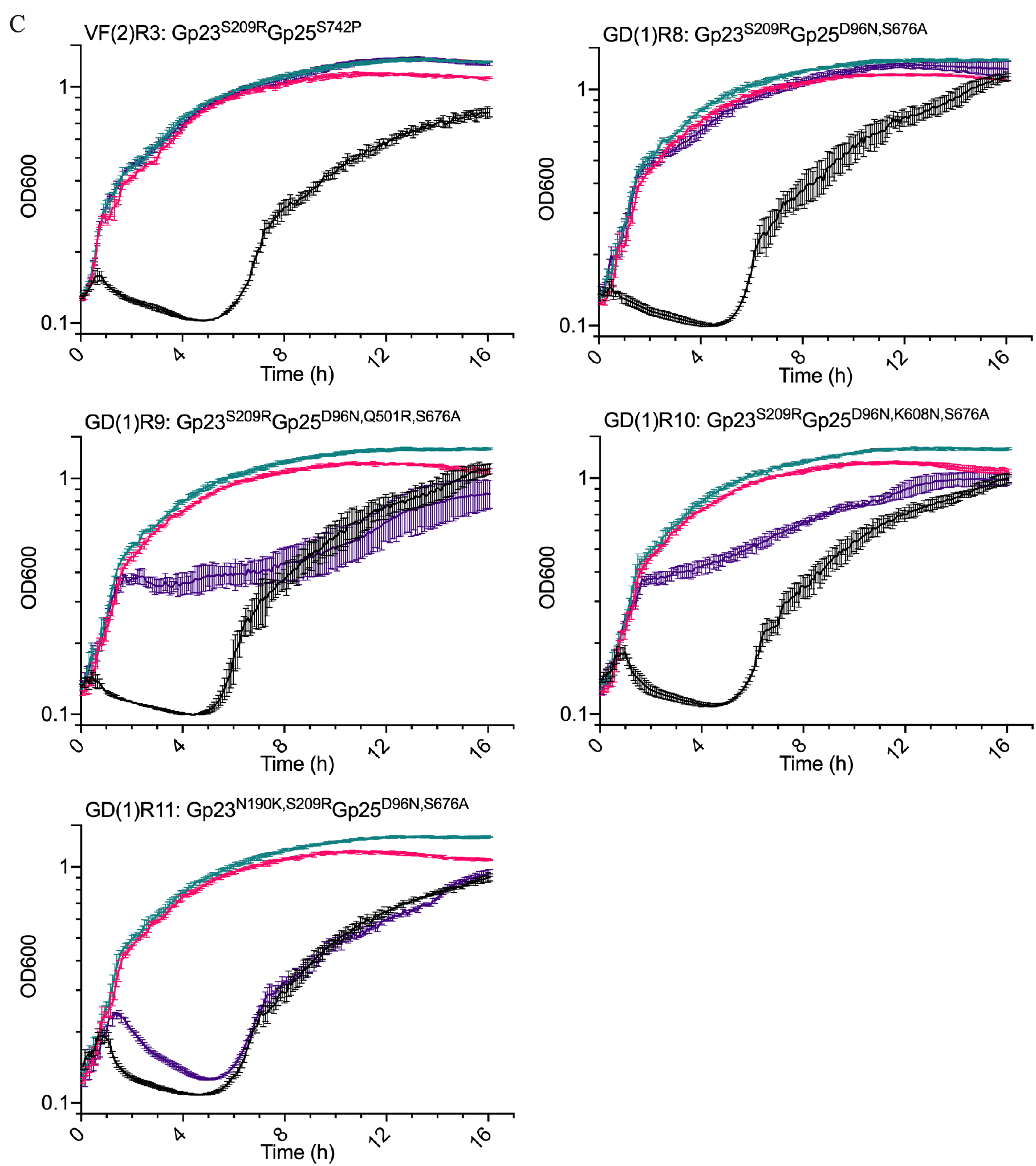
Phage predation killing assays in broth culture. Killing assays were used to evaluate how different ICP2 mutants prey on actively replicating cultures of WT *V. cholerae*, Δ*ompU*, OmpU V324F, and OmpU G325D. Early- to mid-exponential growth phase cells were infected at an MOI ∼1. The optical density at 600 nm was measured every 5 min over 16 hrs at 37°C in a BioTek plate reader. Each graph represents the mean of three technical replicates. Error bars represent standard deviation. A no phage control shows that Δ*ompU* has a slight growth defect. **(A)** WT ICP2 can only prey on WT *V. cholerae*. ICP2 host-range mutants with only Gp25 mutations do not effectively kill OmpU V324F or OmpU G325D. **(B)** ICP2 host-range mutants with a secondary Gp23 S188A kill OmpU V324F with varying degrees of efficiency. **(C)** VF(2)R3 does not kill OmpU G325D despite having a secondary Gp23 S209R mutation, corresponding with its lower EOP and turbid plaque morphology on this host. The remaining ICP2 host-range mutants with Gp23 S209R prey on OmpU G325D, but with varying degrees of efficiency.

In contrast, VF(2)R1, GD(1)R1, and GD(1)R2 do not affect the growth of OmpU V324F or OmpU G325D (Figure 6A), correlating with their lack of binding as shown in Figure 5. These host-range mutants did, however, form turbid plaques and had measurable EOPs on these hosts, presumably due to the slower diffusion of host cells in plaque assays that allows even weak binding phages to eventually infect cells.

ICP2 recombinants with secondary Gp23 mutations specifically prey on either OmpU V324F or OmpU G325D, but not both. Correlating with the intermediate levels of binding in Figure 5, secondary Gp23 mutations result in moderate effects on OmpU V324F or OmpU G325D replication (Figure 6B and C). No host-range mutants tested were able to kill both OmpU V324F and G325D. VF(2)R2 (biological replicate of GD(6)R2) and VF(2)) blocks OmpU V324F late exponential phase. GD(1)R3 and GD(1)R7 do not disrupt OmpU G325D growth consistent with their low EOP (Figure 3B) and weak binding (Figure 5B) on this host.

GD(1)R8, GD(1)R9, GD(1)R10, and GD(1)R11 similarly block OmpU G325D late exponential phase. GD(1)R9 and GD(1)R10 infections lead to a much greater delay than GD(1)R8, despite all three phages having significant EOPs on OmpU G325D (Figure 3C, 6C). Both GD(1)R9 and GD(1)R8 also bind OmpU G325D (Figure 5B). VF(2)R3 has no impact on OmpU G325D growth, correlating with its turbid plaque morphology and reduced EOP (Figure 5C). This further reinforces the necessity of the second Gp25 K159E mutation in GD(6).

## Discussion

### A model of ICP2 evolution within humans

During *V. cholerae* infection in people, the presence of ICP2 phage in the intestinal tract imposes selective pressure, resulting in the appearance of phage escape mutants with mutations in the OmpU receptor. In this scenario, we hypothesize that ICP2 *gp25* mutations are initially selected for increased generalized binding to several OmpU mutants. This generalized binding allows for minimal infection of some OmpU mutant hosts, expanding these phage mutants in the population. Further selection for secondary *gp23* mutations allows for more efficient infection of specific OmpU alleles but comes with the risk of a limited host-range. For example, phages with Gp23 S188A mutations have a narrow host-range and only five of the nine mutants gained an appreciable ability to plaque on one to two additional OmpU alleles. Phages with Gp23 S209R mutations have a wider host-range, with ten of 12 mutants gaining the ability to plaque on three or four additional OmpU alleles.

*V. cholerae* forces ICP2 to generate a variety of different *gp25* and *gp23* mutations, while tempering fitness costs to itself by maintaining a population of functional OmpU alleles (8). This model parallels the one described by DeSordi et al. (25) in which phages infect “intermediate” hosts in the microbiome while evolving towards an expanded host-range. Initial ICP2 host-range mutants with *gp25* mutations are in the process of “jumping” between different OmpU alleles as intermediate hosts. Further studies need to be done to determine if this selective process involves infection processes downstream of OmpU binding, such as DNA injection or virion assembly.

### A model of ICP2 tail fiber structure

Speculation into ICP2 tail fiber structure can be made based on the step-wise evolution of Gp25 and Gp23 and current literature on phage tail fibers and receptor binding proteins (RBPs). Although morphologically in the *Podoviridae* family (7), ICP2 Gp23, Gp24, and Gp25 show surprising similarity to T-even tail fibers in the *Myoviridae* family. Gp25 contains putative collagen-like domains that likely form a triple helical structure, as seen in many other phage tail fibers (26, 27). Gp25 also has a tail tube attachment domain with homology to several other podoviruses (23). Homology analysis via Phyre2 (28) and Hardies et al. (23) revealed that Gp24 has regions very similar to P5 of T4 Gp34 and T4 Gp36 (29). P5 Gp34 is one domain of the T4 long tail fiber closest to the phage baseplate. T4 Gp35 and Gp36 make up the “knee” of the T4 tail long tail fiber (29). None of our ICP2 host-range mutants have Gp24 mutations. Gp23 has both structural (23) and functional homology to T-even adhesins that modulate receptor specificity, such as S16 Gp38 and T4 Gp37 and Gp38 (29-31), suggesting it is the RBP for ICP2.

We hypothesize that *gp25* encodes the long portion of the tail fiber attached to the ICP2 tail tube. Gp25 host-range mutations possibly allow for more permissive receptor binding by Gp23 by altering tail fiber conformation and therefore adsorption. Furthemore Gp25 and Gp23 may interact with independent portions of OmpU or receptors, one of which must be OmpU, altogether. Gp24 makes up the lower long “shin” of a tail fiber and is then distally bound by Gp23, which lies at the tip of each tail fiber. Biochemical and structural studies will be needed to verify this ICP2 tail fiber model. ICP2 Gp14 also has tail fiber homology (23), and we often find missense mutations in this gene, but without selective pressures other than maintaining viability during storage at 4°C (Table S1). Gp14 mutations do not impact Gp23 and Gp25 interactions with OmpU.

### Application

We have yet to find a secondary receptor for ICP2 and the experiments in this study show that ICP2 is dependent on OmpU as its primary receptor. OmpU is a key *V. cholerae* virulence factor (8, 32), limiting its ability to mutate without fitness costs. Our lab has previously shown that a phage cocktail containing ICP1, ICP2, and ICP3 can be used as prophylaxis in two animal models of *V. cholerae* infection (33). In this therapeutic context, these arms-race limitations help ensure both the specificity and efficacy of ICP2.

## Materials and Methods

### Strain construction

A mutator strain of *V. cholerae*, designated AC6727 (Table 1), was constructed by moving marked deletions of *mutS* and the K139 prophage into the rough mutant, AC4653, by natural transformation (34). Genomic DNA was purified from strain AC5218 and transformed into AC4653 with selection for spectinomycin resistance (Sp^R^). Next, a K139-*att* marked deletion was constructed using splicing-by-overlap-extension (SOE) PCR (2), and tranformed into the *wbeL mutS* double mutant with selection for kanamycin resistance (Km^R^). The SOE PCR primers (listed 5’ to 3’) used were: CCATATAAACAACCTAGCTTCGGC, GCTAATACAACATTGAGCCTTGGTG, GGTTCTCTCGCGTTTTACCCCCACCTTTATC, GGGTAAAACGCGAGAGAACCGGGGCTATTTG, CCAGGCTTTACACTTTATGCTTCC, CCCGTCCTAAAACAATTCATCCAG, GGAAGCATAAAGTGTAAAGCCTGGGCGTTTTACCCCCACCTTTATC, and CTGGATGAATTGTTTTAGGACGGGGAGAGAACCGGGGCTATTTG. Strains, phages, and plasmids are listed in Table 1.

**Table 1.** Bacterial strains, phages, and plasmids

| Table 1. Bacterial strains, phages, and plasmids |  |  |
| --- | --- | --- |
| Strain, plasmid, or phage | Description | Source |
| WT | *Vibrio cholerae* O1 El Tor Ogawa E7946 (Sm<sup>R</sup>) | (39) |
| AC2846 | E7946 $\Delta ompU$ | (40) |
| AC6034 | E7946 $\Delta K139$ prophage | Lab collection |
| KS714 | E7946 $\Delta lacZ::Kan^R$, OmpU A182_T193dup | This study |
| KS769 | E7946 $\Delta lacZ::Kan^R$, OmpU V324F | This study |
| KS784 | E7946 $\Delta lacZ::Kan^R$, OmpU A196_Y198dup | This study |
| KS785 | E7946 $\Delta lacZ::Spec^R$, OmpU A195T | This study |
| KS823 | E7946 $\Delta lacZ::Spec^R$, OmpU L319R | (8) |
| KS824 | E7946 $\Delta lacZ::Spec^R$, OmpU S329L | (8) |
| KS825 | E7946 $\Delta lacZ::Spec^R$, OmpU N158Y | (8) |
| KS745 | E7946 OmpU G325D | (8) |
| KS667 | E7946 OmpU A196_Y198dup | (8) |
| KS658 | E7946 OmpU A195T | (8) |
| KS672 | E7946 OmpU V324F | (8) |
| KS822 | E7946 OmpU A182_T193dup, $\Delta lacZ::Spec^R$ | (8) |
| AC4653 | E7469 $\Delta wbeL$ (Sm<sup>R</sup>) | (41) |
| AC5218 | E7469 $\Delta mutS::aad9$ (Sm<sup>R</sup>, Sp<sup>R</sup>) | (24) |
| AC6727 | E7946 $\Delta wbeL \Delta K139-att::aph \Delta mutS::aad9$<br>(Sm<sup>R</sup>, Km<sup>R</sup>, Sp<sup>R</sup>) | This study |
| AC5981 | *E. coli* TG1 | Lab collection |
| AC89 | *E. coli* Sm10λpir | Lab collection |
| pDL1201 | Plasmid backbone for recombination (Amp<sup>R</sup>) | Lab collection |
| pDL1201_gp23-Kan-25 | pDL1201_gp23-neo-gp25 | This study |
| pDL1201_23_GD(6) | pDL1201_gp23(209R)-neo-gp25 | This study |
| pDL1201_23_VF(2) | pDL1201_gp23(S188A)-neo-gp25 | This study |
| pDL1201_25_GD(1) | pDL1201_gp23-neo-gp25(D96N, S676A) | This study |
| pDL1201_25_GD(6) | pDL1201_gp23-neo-gp25(K159E and S742P) | This study |
| pDL1201_25_VF(2) | pDL1201_gp23-neo-gp25(S742P) | This study |
| ICP2 | ICP2_2013_A_Haiti (<u>NC_024791.1</u>) | (8) |

### Bacteriophage isolation and propagation

ICP2_2013_A_Haiti (ICP2) and derivatives were propagated on early exponential growth phase wild-type (WT) *V. cholerae* E7946 (35) growing in Luria Bertani Miller (LB) broth supplemented with streptomycin (100 μg/ ml) at 37°C with aeration. After 3-4 hrs the lysates were centrifuged at 7197 × *g* for 5 min at room temperature (RT) to pellet remaining cells and debris. The supernatants were filtered through 0.22 μm filters and stored as high-titer stocks at 4°C. Stocks were passaged via liquid culture no more than three times to prevent the selection and accumulation of possible mutants. ICP2 stocks were also regularly whole-genome sequenced to authenticate.

Bacteriophage stocks were titered by plaque assay on WT *V. cholerae* within three weeks of being used in experiments. 10 μl of 10-fold serial dilutions of each stock were spotted onto LB 0.3% soft agarose overlays containing approximately 5 × 10^7^ CFU of WT cells and then dribbled across by tilting the plate. Plates were incubated at 37°C overnight (12-18 hrs).

### Genetic analysis of phage isolates

Phage gDNA was isolated from high-titer phage stocks by first pretreating with DNase I and RNase A to hydrolyze contaminating host DNA and RNA. Bacteriophage gDNA was then isolated via phenol-chloroform extraction, or using the DNEasy Blood & Tissue kit (Qiagen) according to the kit instructions and including the optional Proteinase K treatment. DNA samples were prepared for sequencing on an Illumina HiSeq2500 using the Nextera XT Kit (Illumina). CLC Genomic Workbench (Version 20, Qiagen) was used to map the resulting reads to the ICP2_2013_A_Haiti genome (<u>NC_024791.1</u>), and mutations were detected using basic variant detection (Table S2).

### Selection of ICP2 host-range mutants during intestinal infection

All animal experiments were done in accordance with the rules of the Comparative Medicine Services at Tufts University and the Institutional Animal Care and Use Committee. 3-day old infant rabbits were pre-treated with Cimetidine-HCL (Morton Grove Pharmaceuticals) 3 hrs prior to infection (36) and orally inoculated with 5 × 10^8^ CFU of *V. cholerae* in 2.5% sodium bicarbonate (pH 9). A 10:1 mixture of WT to OmpU mutant was used. 5 × 10^6^ PFU of ICP2 was added to the bacteria immediately before inoculation to limit phage adsorption *ex vivo*. Rabbits were euthanized 12 hrs post-inoculation and cecal fluid was collected by dissection and puncture.

### Selection of ICP host-range mutants *in vitro*

A mutagenized pool of ICP2 was generated by preparing two independent high-titer stocks on the *V. cholerae* mutator strain AC6727. Selection was performed on each ICP2 stock by adding 10^10^ PFU to a 1 L, mid-exponential phase, 37^°^C LB broth culture of *V. cholerae* OmpU A182_T193dup. Infection was allowed to proceed overnight. Free phage were purified from the culture supernatant by PEG precipitation. A second, identical selection was performed using the pool of phage purified from each first round selection. Host-range mutants were screened for by plaque assay on the OmpU A182_T193dup mutant. Clear plaque mutants were isolated after each of the second selections, but not from either first selection. Four plaques from each independent selection were plaque-purified on the OmpU A182_T193dup strain. ATdup(1) was isolated from one infection, while ATdup(2) and ATdup(3) were isolated from a second independent infection.

### Homologous recombination of ICP2 *gp23* and *gp25* mutations

Mutant alleles of *gp23* and *gp25* from ICP2 host-range mutants were amplified via PCR and cloned into an ampicillin-resistant plasmid, pDL1201 (Figure S3) by blunt-end ligation (Blunt/TA Ligase Master Mix, New England Biolabs). Mutations in *gp23* or *gp25* were cloned into separate plasmids. In order to mitigate toxicity in *Escherichia coli* and *V. cholerae*, amplified fragments excluded the start and stop codons of *gp23* and *gp25*, and the intervening gene, *gp24*, was replaced with a kanamycin resistance gene (*neo). Neo* also separated recombination events occurring within *gp23* and *gp25*. Additionally, the *gp23-neo-gp25* fragment was fused to a LacZα complementing fragment and downstream of a tight arabinose promotor. Recombinant plasmids were first transformed into chemically competent TG1, then moved into *E. coli* SM10(λpir) by electroporation, and finally mated into WT *V. cholerae* or the ΔK139 strain, AC6034. Use of a ΔK139 strain was done to prevent contamination of ICP2 stocks with the K139 temperate phage, which can spontaneously undergo prophage activation.

Plasmid-containing *V. cholerae* strains were grown to early-exponential phase in LB supplemented with ampicillin and/or kanamycin (50 or 100 μg/ml), infected with ICP2 and then incubated at 37°C for 4-5 hours with aeration. During phage multiplication, recombination with the resident plasmid alleles of *gp23* or *gp25* could occur. The high-titer phage stocks from these infections contained both WT and recombinant ICP2.

### Host-range and efficiency-of-plaquing (EOP) assays

Stocks of each phage were serially diluted 10-fold and dribbled across (10 μl) or spotted on (5 μl) soft agarose overlays containing either WT *V. cholerae* or an OmpU mutant strain. The plates were incubated at 37°C overnight and PFU were counted. Overlays containing *V. cholerae* Δ*ompU* were included as negative controls representing the absence of plaque formation and to rule out contamination by other phages used in the lab. The EOP of ICP2 host-range mutants was calculated as the ratio of PFU on an OmpU mutant divided by PFU on WT. EOPs below 1 indicate that an ICP2 mutant cannot infect a mutant OmpU strain as well as it can infect WT. EOPs were averaged from 2-5 replicate infections. Host-range and EOP assays were scanned to better visualize small and/or turbid plaques (EPSON Scan Version 3.25A).

### Phage binding assays

Mid-exponential phase cultures of WT *V. cholerae* and OmpU receptor mutant strains were washed and resuspended in LB broth. Each strain was diluted to OD_600_ = 0.1 in 200 μl (∼2 × 10^7^ CFU) in PCR tubes (USA Scientific, 1402-4700). Three replicates of each strain were heat-killed at 51°C for 12 min and cooled to 37°C for 2 min in a thermocycler and then cooled to RT before adsorption. Examination using a phase-contrast microscope showed that the heating process did not lyse or disrupt the morphology of the cells. Bacteriophage were added to heat-killed cells at a multiplicity of infection (MOI) of 0.1 (∼2 × 10^6^ PFU). After the addition of phage and mixing, 90 μl were immediately removed and then serially diluted in LB broth to generate a set of samples representing t = 0 hrs of binding. Each dilution series was dribble plated on a soft agarose overlay of WT *V. cholerae*. The remainder of the samples were incubated at RT for 24 hour. After incubation these samples were serially diluted and dribble plated as above. Plates were incubated at 37°C overnight. Binding efficiency was determined as the ratio of PFU at t = 24 hrs to PFU at t = 0 hrs.

Before addition to heat-killed cells, phage stocks with titers below 5 × 10^8^ PFU/ml were extracted with 1-octanol to remove LPS (37, 38) and preheated for 1 hr at 37°C to mitigate discrepancies caused by phage disaggregation at RT. This was not necessary for higher titer phage stocks that required 1:10 or 1:100 dilution before addition to heat-killed cells. 1-octanol extraction and preheating did not affect phage predation (Figure S2).

### Phage predation killing assays

In a 96-well plate, 100 μl of washed mid-exponential phase *V. cholerae* cells diluted to an OD_600_ = 0.2 were infected with 2 × 10^7^ PFU (MOI = 1) in 100 μl of LB broth supplemented with 100 μg/ml of streptomycin (total volume per well = 200 μl). No-phage controls were included to account for growth differences between the host strains. Infections were done in technical triplicate and incubated at 37°C, with shaking at 205 rpm in a plate reader (BioTeK, Synergy H1). OD_600_ was measured every 5 min over 16 hours. This assay was conducted twice for phage mutants without a genetic replicate (Figure S1).

## Acknowledgments

This work was supported by NIH grants GM007310 (ANWL), AI055058 (AC), and AI147658 (AC). AC and MY are associated with and have a financial interest in PhagePro Inc. which is developing a phage cocktail product for prevention of cholera.

