## Supplemental Figure for "A Tail Fiber Protein and a Receptor-Binding Protein Mediate ICP2 Bacteriophage Interactions with *Vibrio cholerae* OmpU"

### Supplemental Figures

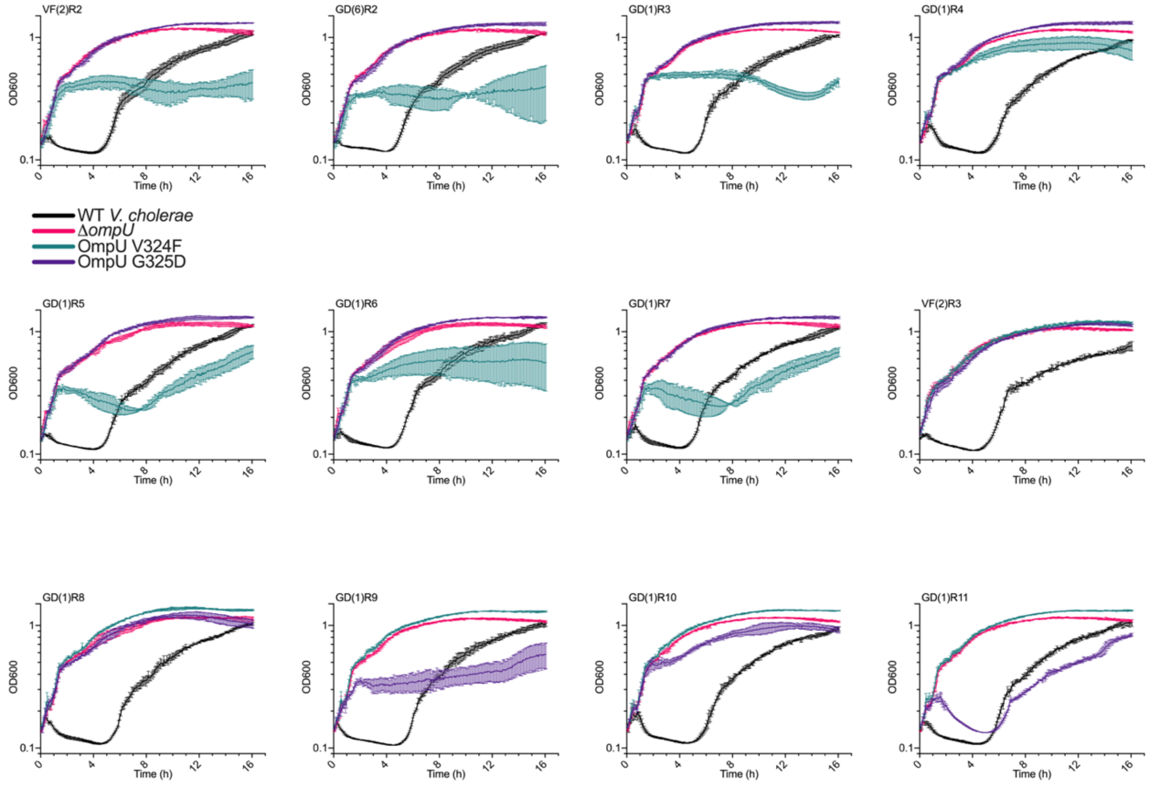


**FIG S1.** Phage predation killing assays were repeated for ICP2 host range mutants that do not have a biological duplicate. Each graph represents the mean of three technical replicates. Error bars show standard deviation. WT ICP2 and VF(2)R1 are not included here because these infections were repeated in Figure S2.

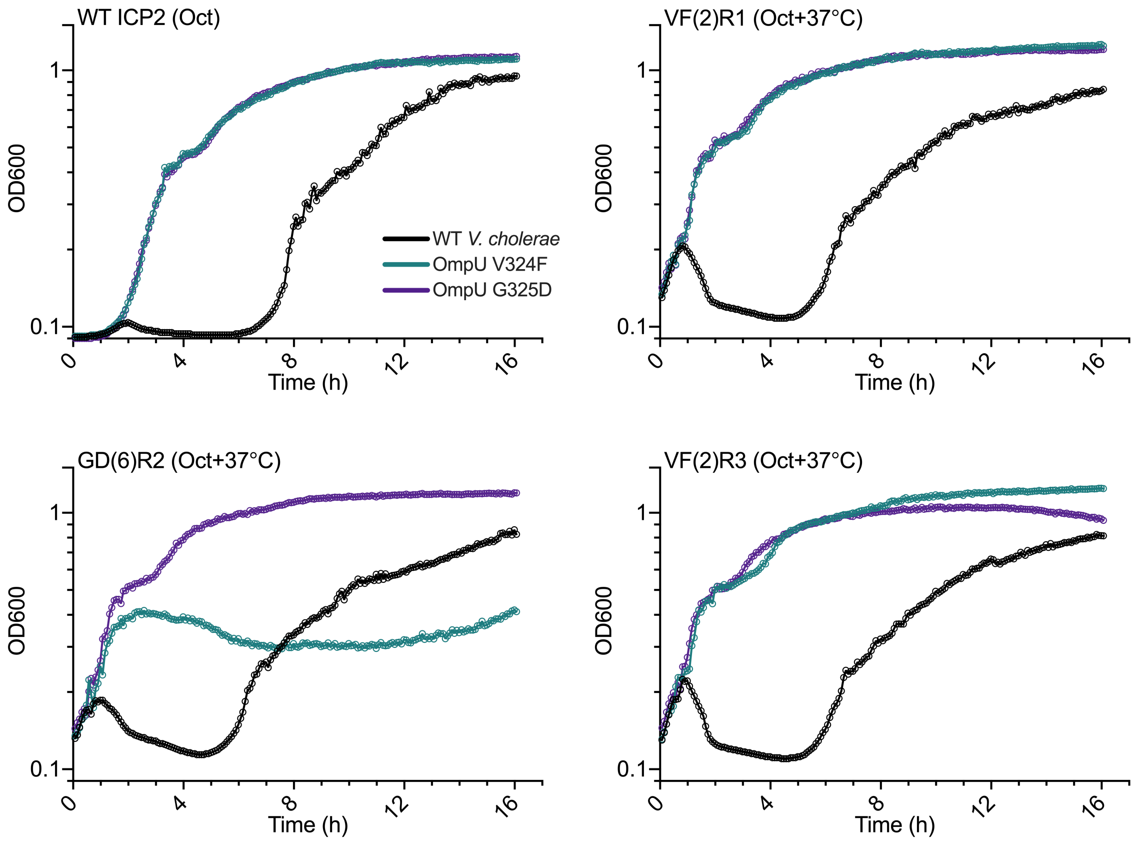


**FIG S2.** Treatment with 1-octanol and/or preheating to 37°C for 1 hr does not greatly affect how WT ICP2, VF(2)R1, GD(6)R2, and VF(2)R3 kill WT *V. cholerae*, OmpU V324F, and OmpU G325D. Early exponential phase cells were infected at MOI ~1. The WT ICP2 stock had a lower titer so the infections were started at a lower cell density. $OD_{600}$ was measured as in the phage predation killing assays. Each set of points represents one replicate.

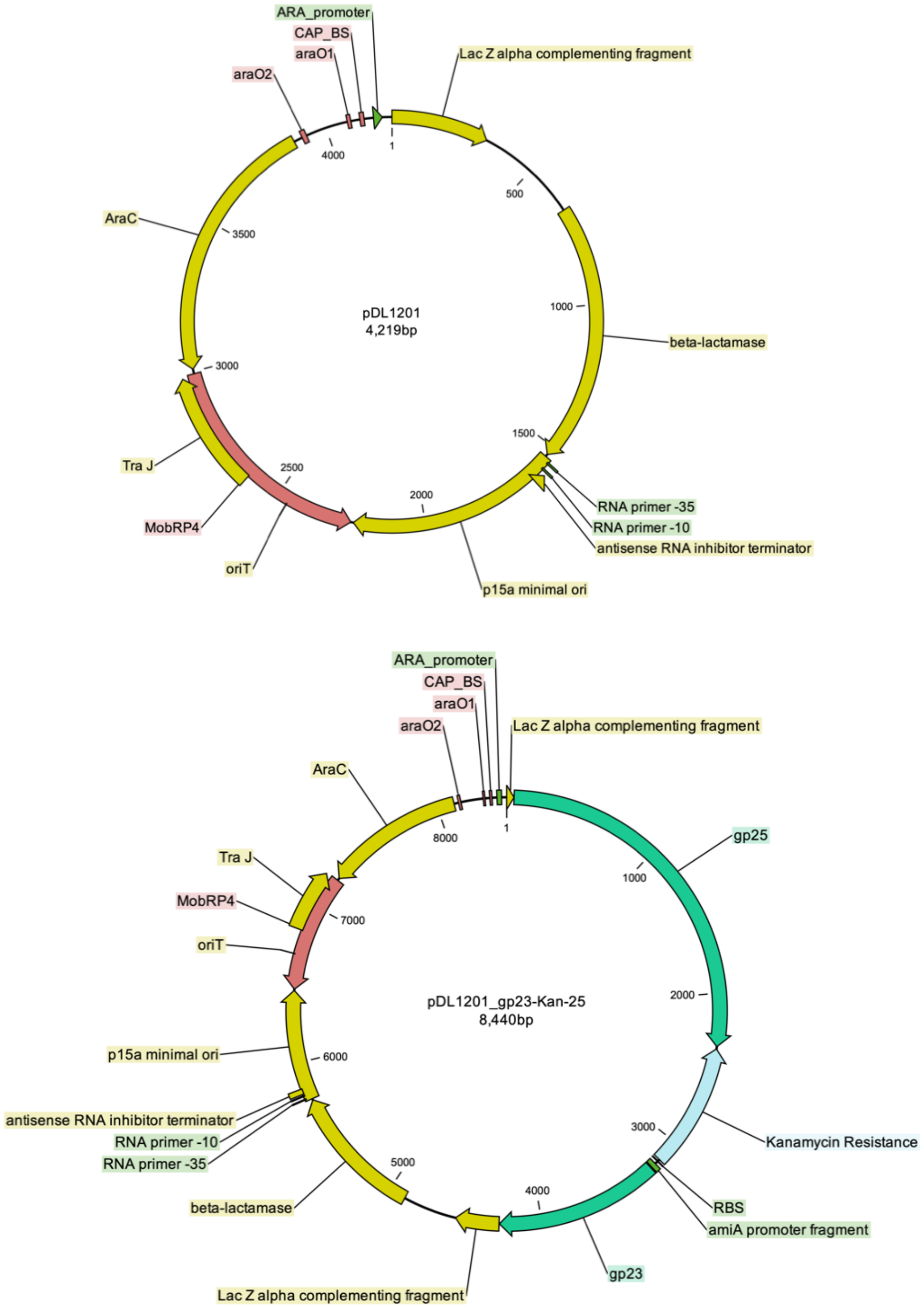


**FIG S3.** Plasmid maps of pDL1201 and pDL1201_gp23-Kan-25. pDL1201encodes a gene for ampicillin resistance, as well as the machinery for bacterial mating. A p15a origin of replication (*ori*) maintains a copy number ~10. While the *gp23-neo-gp25* fragment is presented in one orientation here, the cloning process allowed for ligation in either direction and did not select for one over the other. pDL1201 sequence available upon request.
